# Morphometric analysis of spread platelets identifies integrin α_IIb_β_3_-specific contractile phenotype

**DOI:** 10.1101/264804

**Authors:** Sebastian Lickert, Simona Sorrentino, Jan-Dirk Studt, Ohad Medalia, Viola Vogel, Ingmar Schoen

**Affiliations:** Laboratory of Applied Mechanobiology, Department of Health Sciences and Technology, ETH Zurich, Vladimir-Prelog-Weg 4, 8093 Zurich, Switzerland.; Department of Biochemistry, University of Zurich, Winterthurerstr. 190, 8057 Zurich, Switzerland; Divison of Hematology, University Hospital Zurich, Rämistrasse 100, 8091 Zurich, Switzerland; Department of Life Sciences and the National Institute for Biotechnology in the Negev, Ben-Gurion University, 84105 Beer-Sheva, Israel; current address: Department of Molecular and Cellular Therapeutics and Irish Centre for Vascular Biology, Royal College of Surgeons in Ireland, 123 St Stephen’s Green, Dublin 2, Ireland

## Abstract

Haemostatic platelet function is intimately linked to cellular mechanics and cytoskeletal morphology. How cytoskeletal reorganizations give rise to a highly contractile phenotype that is necessary for clot contraction remains poorly understood. To elucidate this process *in vitro*, we developed a morphometric screen to quantify the spatial organization of actin fibres and vinculin adhesion sites in single spread platelets. Platelets from healthy donors predominantly adopted a bipolar morphology on fibrinogen and fibronectin, whereas distinguishable, more isotropic phenotypes on collagen type I or laminin. Specific integrin α_IIb_β_3_ inhibitors induced an isotropic cytoskeletal organization in a dose-dependent manner. The same trend was observed with decreasing matrix stiffness. Circular F-actin arrangements in platelets from a patient with type II Glanzmann thrombasthenia (GT) were consistent with the residual activity of a small number of α_IIb_β_3_ integrins. Cytoskeletal morphologies *in vitro* can thus inform about platelet adhesion receptor identity and functionality, and integrin α_IIb_β_3_ mechanotransduction fundamentally determines the adoption of a highly contractile bipolar phenotype. Super-resolution microscopy and electron microscopies further confirmed the stress fibre-like contractile actin architecture. For the first time, our assay allows the unbiased and quantitative assessment of platelet morphologies and could help to identify defective platelet contractility contributing to elusive bleeding phenotypes.

## INTRODUCTION

Mechanical platelet functions during thrombosis comprise primary adhesion at injured vessel walls, secondary aggregation, and clot retraction. Each process involves different adhesion receptors and activation pathways but all three require active actomyosin contractile forces. The biophysical mechanisms by which the engagement of different receptors and associated signalling events determine the cytoskeletal architecture associated with the specific platelet sub-phenotypes for adhesion or aggregation, respectively, remain poorly understood. Integrin α_IIb_β_3_-mediated attachment to fibrinogen (FG) results in high single platelet actomyosin contractile forces in the range of 15-35 nN^1–3^. How platelets achieve a similar contraction efficiency as myoblasts^1^ with their highly aligned sarcomeres is unclear. As platelet adhesion and aggregation pose different mechanical requirements, it could be suspected that different cytoskeletal morphologies mediate these different tasks.

The morphology of spreading platelets has been extensively studied *in vitro.* Platelet spreading on glass proceeds fast (1-2 min) and independent of adhesion protein identity^4–6^. Further spreading (3-10 min) on FG or fibronectin (FN) relies on integrin α_IIb_β_3_^4,7^, involves the formation of tight cell-substrate contacts^4^ that are enriched with talin^8,9^, vinculin^8–10^ and Pdlim7^11^, and requires actin remodeling^9^. Fully spread platelets on FG have parallel, triangular, or circular F-actin bundles^9^ that contain myosin, tropomyosin and α-actinin in patches^8,12^, thereby resembling essential features of contractile stress fibres^13^. F-actin arrangements depend on density^14^ and immobilization^15^ of FG and are mediated by integrin signalling pathways involving FAK^15^, Src kinase or Rac^14^. Despite this evidence for a role of integrin outside-in signalling for F-actin cytoskeletal organization, several open questions remain. Are these cytoskeletal arrangements specific for integrin α_IIb_β_3_? Do they contain common signatures that correlate with integrin signalling or mechanotransduction? How do variable integrin α_IIb_β_3_ surface expression levels^16^ or mutations that are associated with Glanzmann thrombasthenia (GT) affect cytoskeletal organization and the capability to develop a contractile phenotype?

A recent study^3^ revealed a potential link between reduced platelet contractility and certain bleeding phenotypes. Since platelet aggregometry tests of these patients were normal, a contractility test could fill a ‘blind spot’ for testing of biomechanical platelet functions. Inhibition of different platelet adhesion receptors resulted in reduced contractility but also distinctly different cytoskeletal morphologies^17^. We thus hypothesize that microscopy images contain valuable information about platelet contractile functions. Until now, platelet cytoskeletal morphologies have been reported without further statistical evaluation^7–10,12^. A systematic high-content screening of platelet morphology, as routinely performed for drug discovery with other adherent cells^18^, is lacking but necessary to establish robust morphological structure-function relationships. We here use confocal fluorescence microscopy to address this relation under well-defined experimental settings. Spreading of washed platelets *in vitro* on ligand-coated surfaces and weak coagulant conditions (5 μM ADP) were chosen to reduce variability. This setting was specifically designed to interrogate the FG – integrin α_IIb_β_3_ – actomyosin interplay which is essential for platelet aggregation. Advanced image analysis reliably identified platelet subpopulations and revealed the predominance of platelets with highly aligned actin cytoskeleton. This morphological phenotype was exclusively linked to α_IIb_β_3_ integrins, depended on their number, clustering, and outside-in signalling capabilities, and was lost on soft matrices or in platelets from a type II GT patient. Platelet cytoskeletal textures thus might serve as biomarkers for haemostatic or defective thrombus formation.

## RESULTS

### Morphological phenotyping of healthy human platelets on fibrinogen

We first imaged human platelets with a cell-permeable F-actin stain^19^ (Supplementary Movie S1 and Supplementary Fig. S1) to determine a seeding time that yields reproducible cytoskeletal morphologies. When platelets touched the FG-coated surface, filopodia and a faint actin ring at the cell periphery appeared. Filament moved radially outwards from the actin-rich centre and bundled. This transient remodelling lasted 3-5 minutes and resulted in strong actin bundles lining a central void region. The timescale of actin remodelling agreed well with dynamics of initial single platelet contraction^1^ and adhesion site formation^4,20^. The stable final arrangement justified the usage of fixed samples for better image quality and higher throughput.

F-actin (Fig. 1a) and vinculin (Fig. 1b) in fully spread platelets showed a wide range of cytoskeletal patterns including bipolar, triangular, star or ring shapes, as reported previously^8,9,12,21^. Despite their variable shape and size, most morphologies were characterized by strong F-actin bundles (Fig. 1a) anchored at pronounced vinculin-containing adhesion sites (Fig. 1b). To systematically analyse these morphologies, aa automated single cell analysis was developed (see Supplementary Text and Supplementary Figs. S2-S4). Platelets’ size and shape, the degree of actin fibre alignment^22^ (from isotropic 0, to completely aligned 1) and their radial order (from circumferential 0, to radial +1) were extracted from images of F-actin (Fig. 1c). The spreading area of platelets from healthy donors ranged from 10 to 60 μm^2^, with a mean at 31 μm^2^ (Fig. 1d). Platelets with areas larger than 20 μm^2^ were regarded as fully spread and subjected to further analysis. Despite their rather circular shape (Fig. 1e), most platelets had a strongly polarized F-actin cytoskeleton with a mean fibre alignment of 0.50 (Fig. 1f) which weakly depended on cell elongation (Fig. S5). No clear preference for radial or circumferential actin fibres was observed (Fig. 1g). The circumferential distribution of vinculin adhesion sites (Fig. 1h) revealed a predominance of bipolar (II, 65%) over triangular (III, 19%) and isotropic (I, 13%) adhesion morphologies (Fig. 1i). Cytoskeletal order and adhesion site distribution were correlated, with aligned and rather star-shaped F-actin for the bipolar morphology, and more isotropic and ring-like F-actin organization for the triangular and isotropic morphologies (Supplementary Fig. S4c+d). This contour plot can thus be regarded as a valid representation of overall adhesion morphologies in the whole platelet population. A representative cell (Fig. 1j) picked from its maximum exemplifies the predominant adhesion phenotype. We conclude that the predominant cytoskeletal architecture of spread platelets on FG in healthy donors was characterized by highly parallel F-actin fibres associated with peripheral adhesions at both ends.

**Figure 1:**
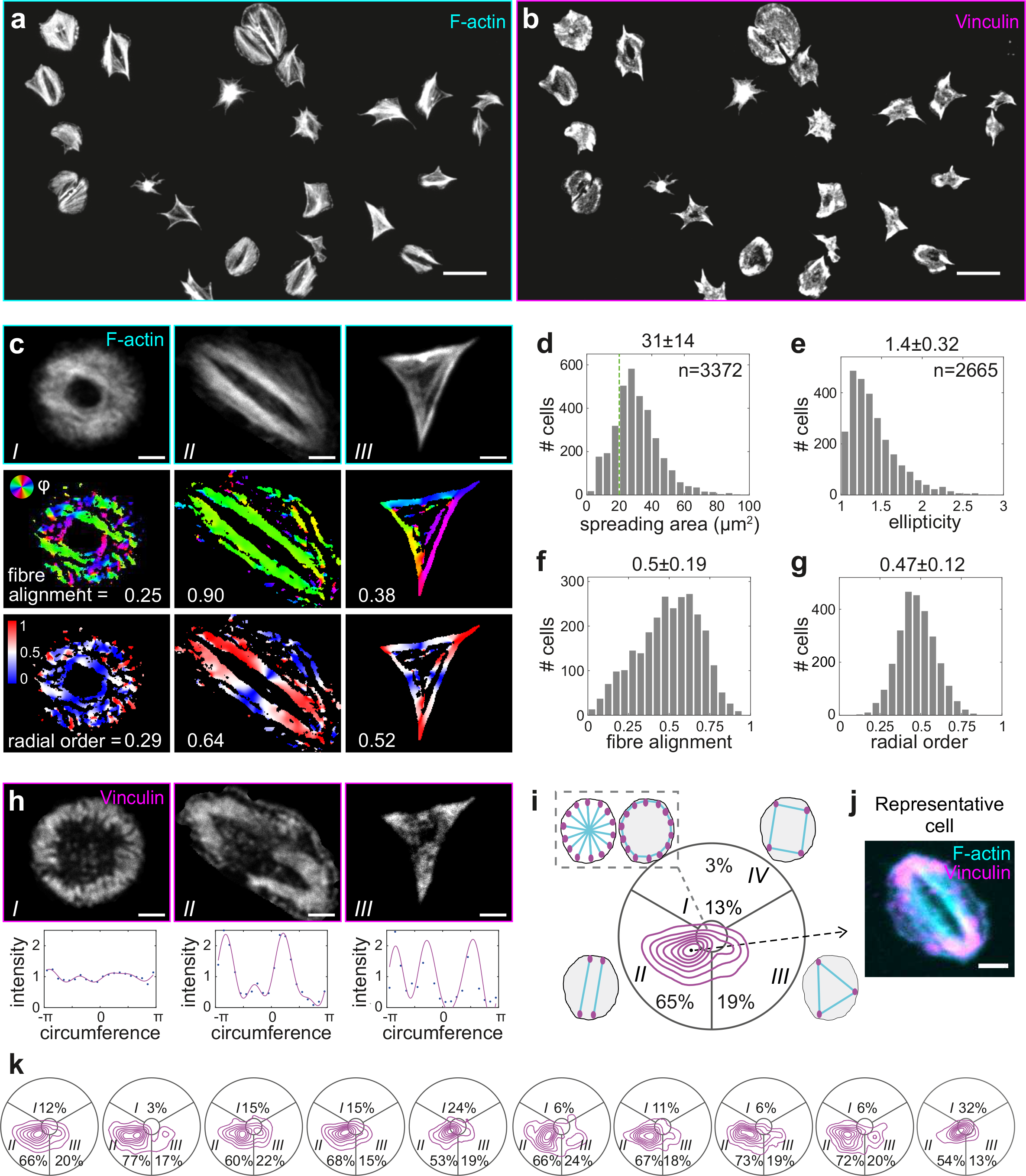
Morphometric analysis of the contractile cytoskeleton in spread platelets reveals pre-dominance of a bipolar phenotype. Representative a) F-actin and b) vinculin confocal micrographs of platelets from a healthy donor after 60 min of spreading on fibrinogen (FG). c) Single cell actin cytoskeletal analysis. Top row: examples of distinctive F-actin patterns. Middle row: Orientation of actin fibres (colour-coded) and derived fibre alignment parameter. 1=perfect alignment, 0=random. Bottom row: Radial order (colour-coded). 1=radial (red), 0=circumferential (blue), 0.5=mixed (white). d) Single cell spreading area. Cells larger than 20 μm^2^ (green dotted line) were further analysed in terms of their e) shape (1: round, >1: elongated), f) fibre alignment, and g) radial order. h) Spatial distribution of adhesion sites. Top row: vinculin stainings, same cells as in c. Bottom row: projected circumferential vinculin intensity profile (dots). A Fourier fit (magenta solid line) was used to extract the principle components up to 4th order (bipolar, triangular, and quadratic arrangements). i) Contour plot of the number of cells with adhesion sites arranged isotropically (middle sector) or in a bipolar (lower left sector), triangular (lower right sector), or quadratic (upper sector) pattern. j) Representative cell for the predominant morphology picked according to the maximum in the vinculin morphology plot (dot in i). Data were pooled from 10 healthy donors with 200-400 platelets per donor. k) Comparison of donor-to-donor variability by vinculin morphology (cf. also Supplementary Fig. S9). Donors comprised 7 males and 3 females with a median age of 31.5 years (range 27-44 years). Scale bars: 10 μm (a,b), 2 μm (c,h,j). A comprehensive description of the image analysis is given in the Supplementary Text and Supplementary Figs. S2-S4.

### Assay validation and physiological variability

We next assessed the repeatability and reproducibility of platelet adhesion morphologies. Independent processing of the same blood sample yielded highly consistent results (Table 1, ‘intra’). The morphology was robust with prolonged incubation up to four hours (Supplementary Fig. S6). In the absence of ADP, platelet spreading area, ellipticity, and radial order were unaffected whereas the fibre alignment was slightly decreased (Supplementary Fig. S7). Storage of the blood sample for 24 hours before analysis had a similar effect (Supplementary Fig. S8). We conclude that platelet adhesion morphology was highly reproducible under the chosen assay settings, i.e. in the presence of ADP and within a few hours after blood withdrawal.

**Table 1:** Assay validation and physiological variability. Given are the mean differences in absolute values or the respective coefficients of variation (CV, in brackets) of morphometrics between different samples for spreading area, ellipticity, fibre alignment, and radial order. The similarity between vinculin morphology plots was quantified by a normalized cross-correlation (‘CC^morph^’). Intra: four independently processed samples from the same blood probe. In-donor: two samples each from repeated withdrawals, n=4 donors. Inter-donors: samples from n=10 healthy donors.

| | Area/$\mu\text{m}^2$ | Ellipticity | Fibre alignment | Radial order | CC<sup>morph</sup> |
| --- | --- | --- | --- | --- | --- |
| intra | 1.30 (0.04) | 0.039 (0.10) | 0.022 (0.04) | 0.007 (0.01) | 0.91 (0.02) |
| in-donor | 3.68 (0.12) | 0.104 (0.25) | 0.016 (0.03) | 0.050 (0.10) | 0.91 (0.02) |
| inter-donor | 3.62 (0.12) | 0.064 (0.16) | 0.069 (0.14) | 0.046 (0.09) | 0.83 (0.10) |

To test the physiological variability of platelets from the same donor, withdrawals were repeated at intervals of more than one week. Resulting CVs were only slightly higher (Table 1, ‘in-donor’) than the assay reproducibility. When assessing variation between donors, the contour plots consistently showed primary bipolar and secondary triangular subpopulations (Fig. 1k). Statistically significant differences between donors were detected in few cases (Supplementary Fig. S9), and CVs were of similar magnitude (Table 1, ‘inter-donor’) as for repeated withdrawals. In conclusion, actin and vinculin morphometrics were more sensitive than spreading area and shape, morphological signatures were donor-specific, yet the bipolar morphologies and strong fibre alignment dominated in all healthy individuals.

### Specificity of the bipolar morphology for ligands of integrin α_IIb_β_3_

To test whether the morphology *in vitro* correlates with the engagement of specific adhesion receptors, we seeded platelets on surfaces coated with FG, fibronectin (FN), laminin (LN), or collagen type 1 (COL1). FG and FN are bound primarily by integrin α_IIb_β_3_ in the context of platelet aggregation, whereas COL1 is recognized by integrin α_2_β_1_ (and GPVI) and LN by integrin α_6_β_1_ in the context of platelet adhesion to the sub-endothelium^23^. Platelets adhered to all surfaces (Fig. 2a). The bipolar phenotype dominated on FG and FN, whereas most platelets on LN and COL1 showed isotropic or disordered adhesion patterns (Fig. 2b). Spreading areas on FG, FN, and LN were comparable (mean 30 μm^2^) but smaller on COL1 (mean 20 μm^2^; Fig. 2d). Platelet shape was similar on FG, FN, and COL1 but rounder on LN (Supplementary Fig. S10a). The F-actin cytoskeleton was strongly aligned on FG and FN (mean >0.5) but strikingly less on LN (mean 0.4) and COL1 (mean 0.35; Fig. 2e). On LN, it was arranged in a pronounced circumferential ring^24,25^ (Fig. 2a,b) as reflected by the lower radial order (mean 0.35; Supplementary Fig. S10b). The spatial distribution of vinculin adhesion sites on FG and FN was similar but substantially different on COL1 and LN (Fig. 2f). In summary, platelets developed the same morphologies on FN and FG, but highly discriminable phenotypes on other adhesion proteins.

**Figure 2:**
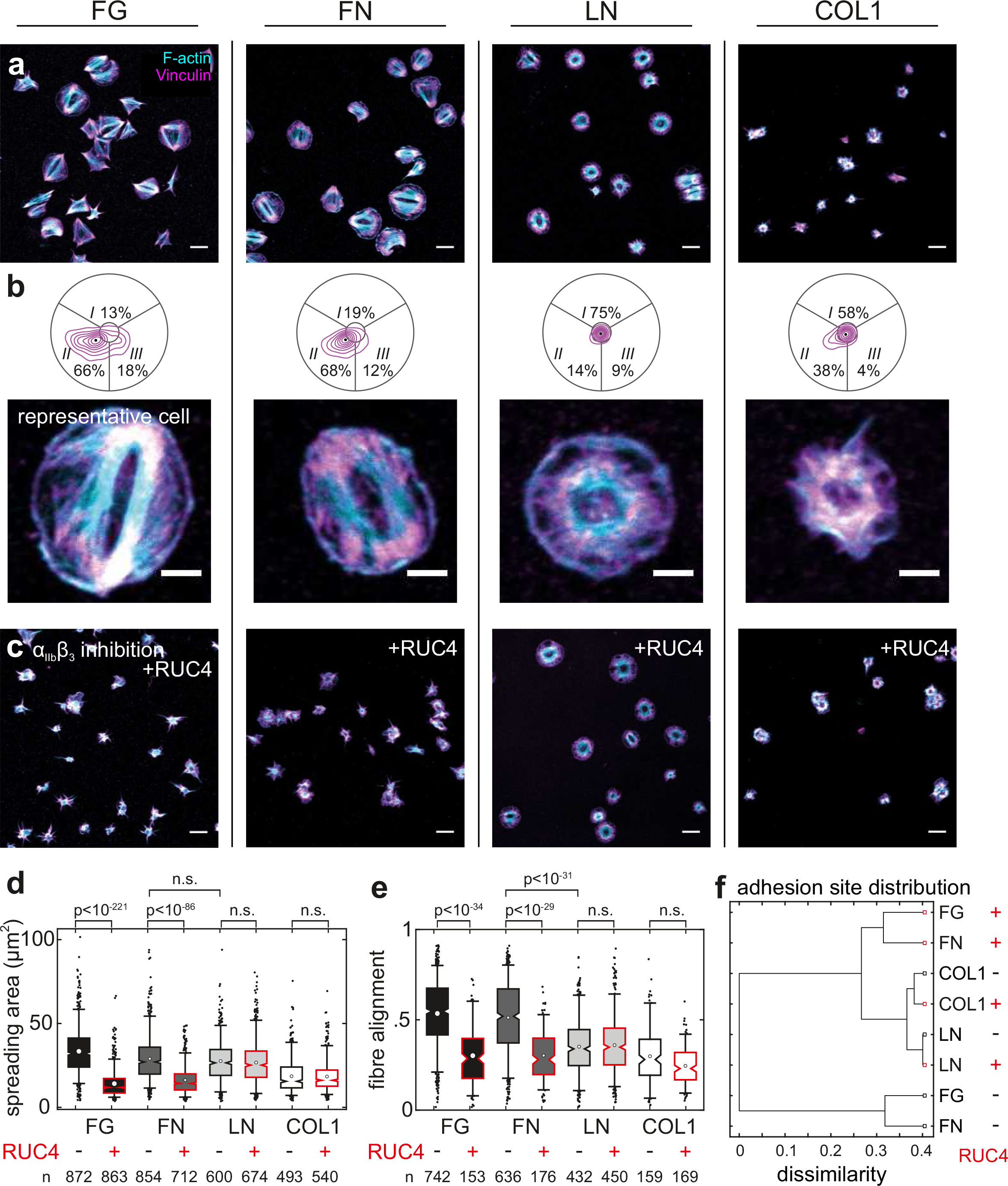
The bipolar phenotype is exclusively observed on ligands of integrin α_IIb_β_3_. a) Overview of healthy platelets after 60 min of spreading on the ligands FG, fibronectin (FN), laminin (LN) and collagen type I (COL1). Cyan: F-actin; magenta: vinculin. b) Spatial distribution of vinculin adhesion sites in platelets and representative cells. c) Spread platelets in the presence of 100 μM of the α_IIb_β_3_-specific inhibitor RUC-4. Quantitative F-actin analysis for d) the spreading area and e) the fibre alignment is shown for each ligand with and without blocking the integrin α_IIb_β_3_ (cf. also Supplementary Fig. S10). f) Classification according to the similarity between vinculin adhesion site distributions. The classification tree clearly shows that morphologies on FG and FN are very similar and heavily affected by inhibiting α_IIb_β_3_ whereas COL1 and LN are very different from this morphology and from each other and not affected by RUC-4. Data were pooled from three healthy male donors (33-44 years). Scale bars: 10 μm (a,c), 2 μm (b).

To determine the contribution of integrin α_IIb_β_3_ to morphologies on different adhesion proteins, we applied the specific inhibitor RUC-4^26^ at saturating (100 μM) concentration. The presence of RUC-4 effectively abolished platelet spreading on FG and FN, whereas on LN or COL1 it had no significant effect on platelet morphologies (Fig. 2c-f). We conclude that the bipolar phenotype is exclusively associated with adhesion ligands that play a role in platelet aggregation and it requires integrin α_IIb_β_3_.

### Sensitivity of the bipolar morphology to integrin α_IIb_β_3_ outside-in signalling

Integrin α_IIb_β_3_ is expressed at ∼80 000 copies per platelet^27,28^, yet the functional importance of this high copy number and its variations remain unclear. To distinguish between the effects of binding strength, clustering and outside-in signalling on platelet morphology, three different approaches were chosen to perturb the number of engaged α_IIb_β_3_ integrins.

First, we performed a dose-response experiment with RUC-4 that blocks access to the binding pocket of integrin α_IIb_β_3_ but does not ‘prime’ it for outside-in signalling^26,29–31^ (Fig. 3a,i). F-actin became gradually less aligned (IC_50_ 1.3 μM, Fig. 3a,iii) with increasing RUC-4 concentrations before the spreading area was affected (IC_50_ 6.3 μM, Fig. 3a,ii). Concomitantly, the population shifted from the bipolar to a more isotropic phenotype (Fig. 3a,iv). The DMSO control was negative (Supplementary Fig. S11). Hence, a high number of functional α_IIb_β_3_ integrins was needed for the bipolar phenotype.

**Figure 3:**
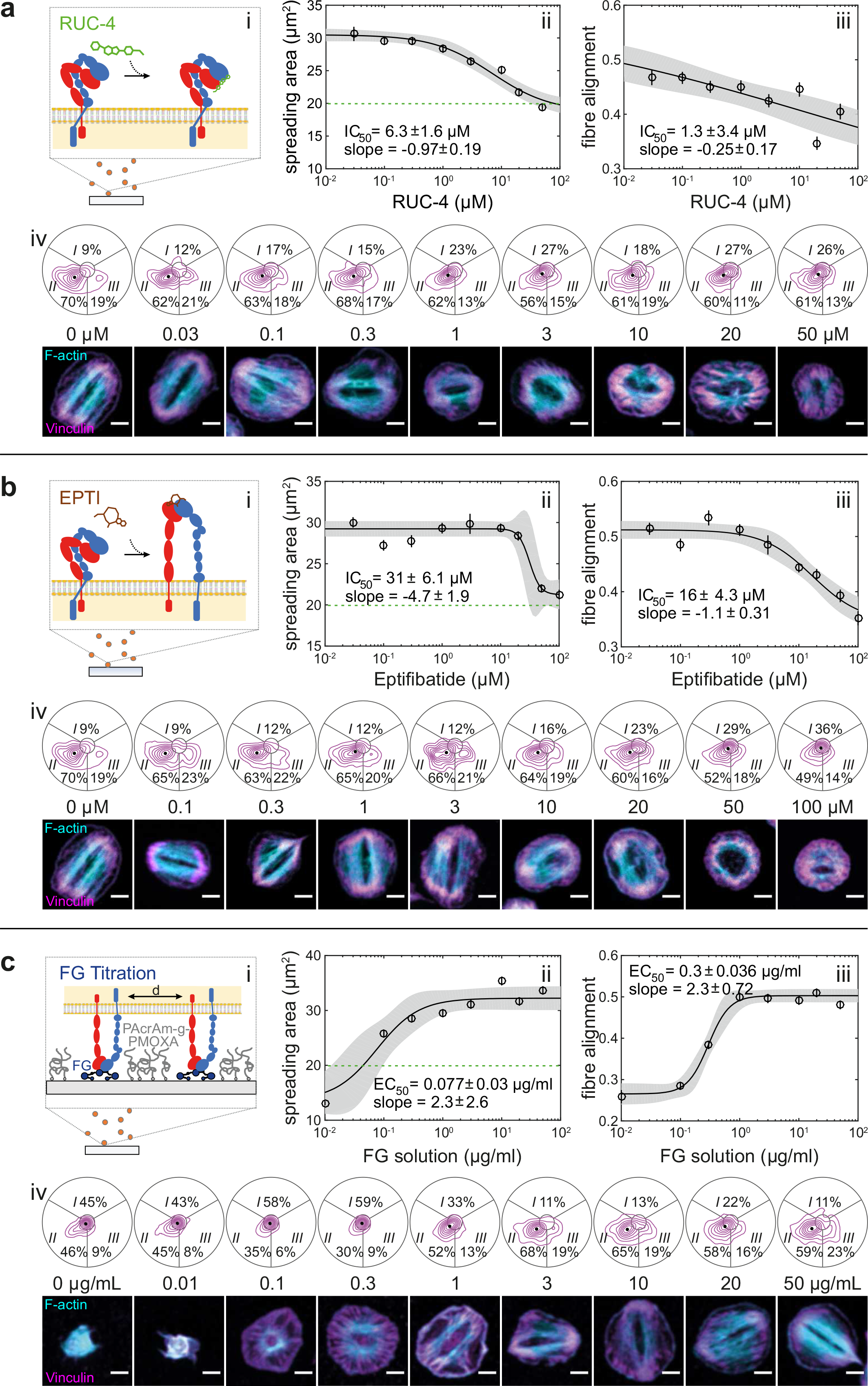
The bipolar morphology depends on integrin α_IIb_β_3_ outside-in signalling. a) Titration series with the non-priming inhibitor RUC-4 that blocks FG binding of α_IIb_β_3_. b) Titration series with Eptifibatide which induces conformational changes in related to outside-in signalling. c) Dilution series of surface-immobilized FG that affects integrin clustering (cf. also Supplementary Figs. S12 and S13). Shown are i) schematic representations, dose-response curves for ii) spreading area and iii) fibre alignment, and iv) contour plots of the spatial distribution of adhesion sites together with representative cells. Solid lines and grey regions are fits to a logistic Hill equation and their 95% confidence intervals, respectively. Data pooled from three male donors (33-44 years) each. Platelets were preincubated for 10 minutes with inhibitors and seeded for 60 minutes in their presence. Scale bars: 2 μm.

Second, we used the α_IIb_β_3_ antagonist eptifibatide (Integrilin) that causes ligand-induced conformational changes related to outside-in signalling^32^ similar to natural ligands like FG^33^ (Fig. 3b,i). F-actin alignment decreased abruptly (IC_50_ 17 μM, Fig. 4b,iii) and the bipolar phenotype disappeared (Fig. 3b,iv) shortly before spreading was inhibited (IC_50_ 31 μM, Fig. 3b,ii). The about 5-7 fold steeper dose-response curves compared with RUC-4 indicate that eptifibatide-induced outside-in signalling might partially compensate for the reduced number of engaged integrins.

**Figure 4:**
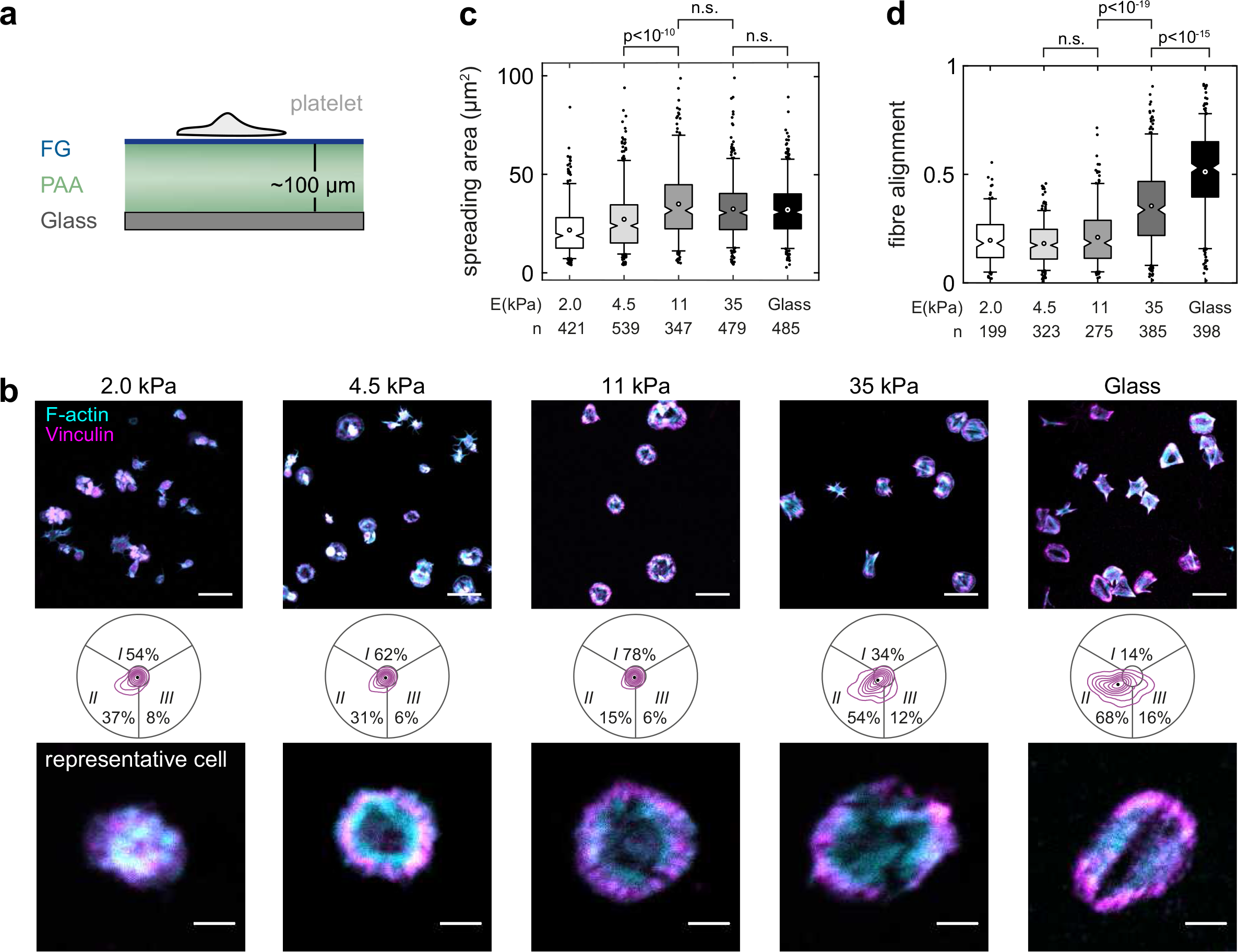
The bipolar morphology is mechanosensitive. a) Platelets were seeded for 60 min on FG that was covalently crosslinked on polyacrylamide (PAA) hydrogels of different stiffness. Schematic not to scale. b) Overview images (top), adhesion morphology plots (middle) and representative cells (bottom) of platelets on 2.0 kPa, 4.5 kPa, 11 kPa, 35 kPa gels and on glass. Scale bars: 10 μm (overview), 2 μm (repesentative cells). c) Cells spread on gels above 4.5 kPa. c) Actin fibre alignment increased above 11 kPa. Data were pooled from one male and two female donors (28-54 years). See also Supplementary Fig. S14.

Third, we varied the density of surface-immobilized FG (Supplementary Fig. S12) which affects integrin clustering, early adhesion stabilization, integrin signalling and platelet spreading^14^ (Fig. 3c,i). Unspecific surface interactions were blocked by a backfill with the non-fouling agent PAcrAm-g-PMOXA^34,35^ (Supplementary Fig. S13). The morphology of platelets changed gradually with decreasing FG surface density. A pronounced ring-like F-actin and vinculin arrangement prevailed at intermediate densities (around 0.2 μg/mL, Fig. 3c,iv) which was accompanied by reduced actin alignment (Fig. 3c,iii) and a loss of the bipolar signature (Fig. 3c,iv). Spreading was suppressed below 0.1 μg/mL (EC50 0.08 μg/mL, Fig. 3c,ii). We conclude that clustering of integrin α_IIb_β_3_ sensitively affected the cytoskeletal organization and was required for the bipolar phenotype.

In summary, these perturbations of integrin α_IIb_β_3_ reveal an important role of functional integrin outside-in signalling and clustering for the bipolar morphology.

### Dependency of the bipolar morphology on mechanosensing through integrin α_IIb_β_3_

Matrix stiffness modulates the assembly and remodeling of integrin junctions^36–38^ and affects cell spreading^39^, cytoskeletal organization^22^, traction forces^39,40^, and platelet activation^41^. To study the effect of integrin α_IIb_β_3_ mechanosensing on cytoskeletal morphology, platelets were seeded on FG-coated polyacrylamide hydrogels with a stiffness between 2-35 kPa (Fig. 4a). Platelets spread on all but the softest gel (Fig. 4b,c), in agreement with previous findings^41^. Like on glass, most platelets on the hardest gel showed a bipolar signature (Fig. 4b) and an aligned actin cytoskeleton (Fig. 4d). Below 11 kPa, the bipolar phenotype was gradually lost and actin organization became more isotropic (Fig. 4b,d). These morphological features resembled those for intermediate RUC-4 concentrations (cf. Fig. 3a) or reduced FG density (cf. Fig. 3c). Please note that the FG density was similar on all gels^42^ (Supplementary Fig. S14). We conclude that an extracellular resistance to traction forces through integrin α_IIb_β_3_ is needed to achieve a strong alignment of the cytoskeleton.

### Ultrastructure of the healthy cytoskeletal morphology on fibrinogen

Although electron and super-resolution microscopes are not yet well suited for a high throughput screen, they here were used to further substantiate our conclusions derived from confocal microscopy. Most microfilaments in SEM images ran in parallel from one side of the cell to the other and fused in-between into larger bundles (Fig. 4a, inset, arrowheads), as previously reported^43,44^. Direct stochastic optical reconstruction microscopy (dSTORM)^45^ of F-actin revealed densely arranged and strongly aligned actin filaments within bundles (Fig. 5b and Supplementary Fig. S15A), whereas F-actin in the cell periphery formed a dendritic network (Fig. 5b, inset) as also obvious from cryo-electron tomograms (cryo-ET; Supplementary Fig. S16a). The good agreement between SEM with cryo-ET images and F-actin dSTORM images indicates that the majority of filamentous structures in SEM images were actin microfilaments.

**Figure 5:**
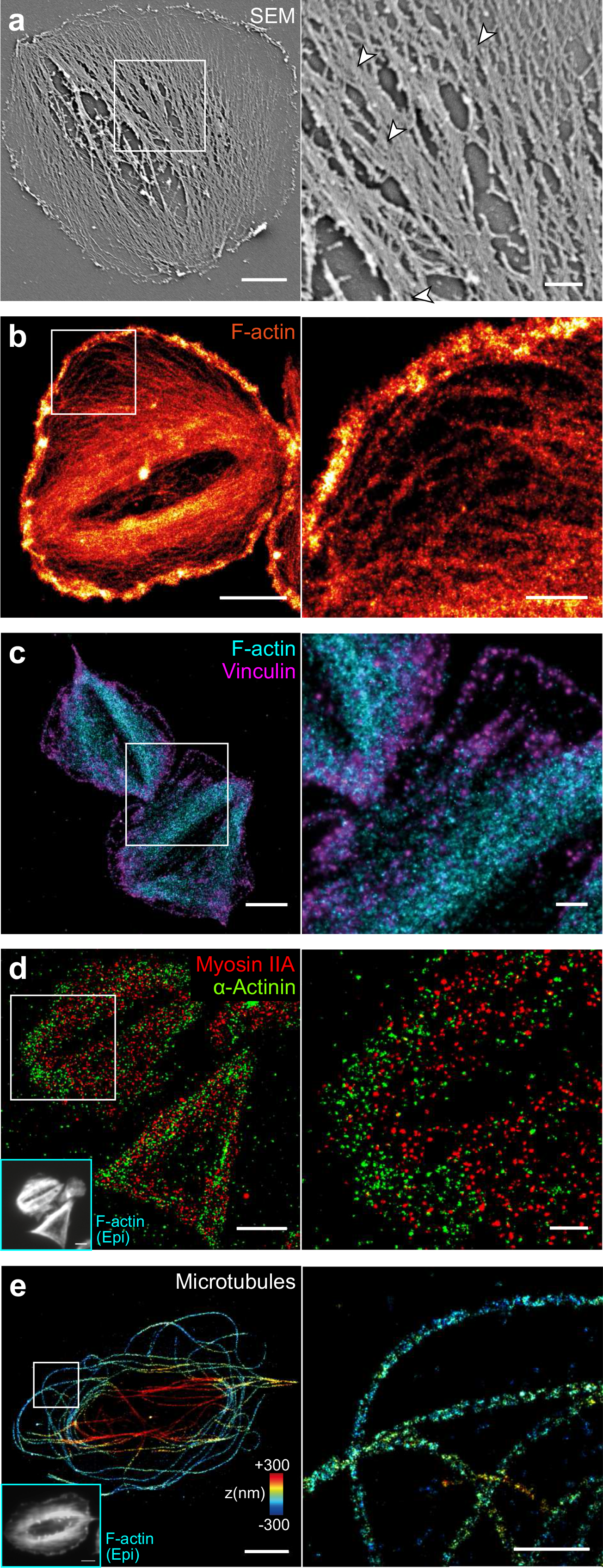
Electron microscopy and superresolution (dSTORM) imaging of the cytoskeletal morphology of healthy human platelets. Platelets were seeded for 60 min on FG on glass, then detergent extracted and fixed. Shown are representative cells (left) and a magnified inset (right). a) Scanning electron microscopy (SEM) image. b) dSTORM image of F-actin. c) Dual-colour dSTORM image of vinculin (magenta) and F-actin (cyan). d) Dual-colour dSTORM image if mysoin IIA (red) and α-actinin (green). Inset: epifluorescence image of F-actin in the same cells. e) 3D dSTORM image of microtubules. The z-position is colour-coded from blue (basal) to red (apical). Individual microtubules are characterized by the rail-track-like projection of their immunolabelled outer shell. Inset: epi fluorescence image of F-actin in the same cell. Scale bars: 2 μm (left column), 500 nm (insets). See also Supplementary Fig. S15 for overview dSTORM images and Supplementary Fig. S16 for cryo-electron tomograms.

We further investigated the composition of actin bundles and their respective adhesion sites. Dual-colour dSTORM images showed string-like vinculin stainings along the ends of actin fibres (Fig. 5c and Supplementary Fig. S15b) similar to observations in focal adhesions of fibroblasts^46^. The motor protein myosin IIA co-localized with actin bundles (Fig. 5d, red) and was phosphorylated at its light chain (Supplementary Fig. S17), confirming that F-actin bundles were contractile. The actin-binding protein α-actinin was localized in adhesion sites and lesser in actin bundles (Fig. 5d, green). Our inability to detect an alternating arrangement of these two latter proteins along F-actin bundles (Fig. 5d, inset; in contrast to a previous report^12^) or bipolar myosin mini-filaments as two hallmark features of stress fibres^13,47,48^ might be partially owed to the poor immunolabeling efficiency for myosin and α-actinin.

As the marginal band dominates the passive mechanical properties of resting platelets^49^, we visualized microtubules by 3D dSTORM (Fig. 5E and Supplementary Fig. S15c). In agreement with previous observations^50,51^, microtubules in fully spread platelets on FG mostly formed uncurled, spaghetti-nest like structures, with few remnants of the marginal band. Some microtubules bridged the granulomere at the apical side, whereas others extended into the lamellipodial region (Fig. 5e, inset). Overall, no clear association of microtubules with actin bundles was apparent.

Taken together, the ultrastructure of human platelets confirms that their active mechanical properties are dominated by F-actin microfilaments which are bundled into largely parallel stress fibre-like structures.

### Change of platelet phenotype in Glanzmann thrombasthenia

Certain mutations in integrin α_IIb_β_3_ cause GT, a rare bleeding disorder of autosomal-recessive inheritance accompanied by defective aggregation and clot retraction^52^. To investigate a possible impact on platelet morphology, we recruited a 54-year old female patient with well characterized GT: light transmission aggregometry in platelet-rich plasma demonstrated an impaired platelet aggregation with all agonists (arachidonic acid, thromboxane receptor agonist U46619, ADP, collagen, epinephrine) except ristocetin. Flow cytometry showed significantly reduced (6-7% of normal) surface expression levels of both α_IIb_ and β_3_. The ITGB3 gene carried a heterozygote small duplication in exon 10 (p.Asn470Ter) and a heterozygote missense-mutation in exon 11 (p.Gly605Asp). The duplication is expected to result in severely reduced integrin β_3_ levels due to pre-mature termination of protein expression. The second mutation affects the same residue as two other described mutations^53,54^ which were associated with an abnormal reduction of surface expressed α_IIb_β_3_ and its constitutive activation (Supplementary Fig. S18).

Platelets from this GT patient spread on FG in the presence of ADP (Supplementary Fig. 6a). After 60 minutes, they were slightly smaller (mean 26 μm^2^) than healthy platelets but significantly rounder in shape (Supplementary Fig. 6b) and showed pronounced concentric F-actin and vinculin stainings (Fig. 6a,c). Consequently, actin fibre alignment (mean 0.26) and radial order (mean 0.29) were significantly reduced (Fig. 6b) and 81% of platelets had an isotropic phenotype (Fig. 6c). Spreading was completely abolished by RUC-4 (Supplementary Fig. S19) which proofs its dependence on α_IIb_β_3_. Results were highly reproducible between separate withdrawals (Supplementary Fig. S20).

SEM images (Fig. 6d) and dSTORM images of F-actin (Fig. 6e and Supplementary Fig. S21a) showed circumferential actin fibres lining the central granulomere but no observable bundling. The dendritic actin network in the lamellipodium was normal (Fig. 6e and Supplementary Fig. S15b). Vinculin adhesion sites were located just outside the F-actin ring (Fig. 6f) with distinct string-like arrangements in the radial direction (Fig. 6f, inset, and Supplementary Fig. S21b). The actin ring zone was positive for pMLC (Supplementary Fig. S22) and thus presumably contractile. Frequent remnants of the microtubule marginal band (Supplementary Fig. S23), reduced spreading in the absence of ADP (Supplementary Fig. S20), and an increased cytoskeletal alignment after prolonged incubation (Supplementary Fig. S24) hint towards an incomplete activation. We conclude that platelets from this GT patient were not able to assemble and stabilize their cytoskeleton towards the healthy bipolar phenotype.

## DISCUSSION

Since reproducibility in the data analysis of cells poses a severe challenge, we introduce an easy-to-implement screening assay for assessing platelet cytoskeletal morphology. It is based on dual-channel (immuno)fluorescence confocal images (Fig. 1) plus automated image analysis (Supplementary Figs. 2-4) and yields highly reproducible metrics (Table 1). In contrast to more complex approaches^18^, the measured parameters reflect features that are directly related to mechanical cell function^22^ and can be visually verified.

Engagement of the platelet integrin α_IIb_β_3_ was necessary for a predominant bipolar organization as shown by ligand selectivity (Fig. 2) and specific blocking (Figs. 2c+d, 3a+b). Its disappearance at sub-saturating concentrations of RUC-4 (Fig. 3a), at reduced FG surface densities (Fig. 3c), or in platelets from a GT patient with reduced α_IIb_β_3_ surface expression levels (Fig. 6) show that a small number of engaged integrins was insufficient to induce cytoskeletal polarization. The late onset of cytoskeletal changes with the ‘priming’ ligand eptifibatide at around 3 μM (∼95% receptor occupancy^55^; Fig. 3b) implies that strong α_IIb_β_3_ outside-in signalling could compensate for small numbers of bound integrins and therefore promote the bipolar phenotype. This view is supported by the finding that low (3 μg/mL) FG densities alter Src, Rho, and Rac signalling^14^ downstream of α_IIb_β_3_. Selective inhibitors against these pathways caused F-actin morphological changes^14^ that resemble the isotropic phenotype seen with integrin inhibitors (Fig. 3a,b)^20^, on low FG densities (Fig. 3c), or on soft hydrogels (Fig. 4b). The higher cytoskeletal ordering in platelets on stiffer matrices (Fig. 4d) agrees with mechanosensing mechanisms described for mesenchymal cells, including stem cells^22^, which rely on integrin mechanotransduction, adhesion maturation, and increased traction forces^37,38^. Mechanosensing through α_IIb_β_3_ thus simultaneously controls the number of engaged and signalling integrins and thereby dictates cytoskeletal morphology. In summary, the bipolar signature can be seen as a ‘morphological fingerprint’ of functional integrin α_IIb_β_3_ outside-in signalling.

Our results have implications for how platelets pull on their environment. The parallel organization of actomyosin filaments in the bipolar phenotype on FG (Figs. 1, 5) enables an efficient force generation and transduction, without the need for an elongated cell shape as in stem cells^22,56^ and sarcomere-like ordering as in cardiomyocytes^57,58^, and supports high single platelet forces^1–3^. This cytoskeletal polarization is expected to result in anisotropic (dipolar) tractions^20^. The rather isotropic force fields measured by traction force microscopy (TFM) of platelets on soft hydrogels^2^ might be explained by the limited spatial resolution (∼1-2 μm) of TFM and reduced cytoskeletal polarization (Fig. 4D) and contractility^41,59^ at 4.5 kPa. The fact that the bipolar phenotype was exclusively observed on FG and FN, but not on COL1 or LN (Fig. 2), suggests that the platelets’ capability to polarize their cytoskeleton might especially be relevant in the context of platelet aggregation, aiding efficient clot retraction.

In conclusion, this study demonstrates the first direct morphometric high-content screening of platelets and the identification of subpopulations that differ in their functional phenotypes. The tight correspondence of platelet cytoskeletal morphology with a mild GT bleeding phenotype (Figure 6) and with the action of aggregation antagonists (Figure 3) suggests that a morphological investigation of platelets might help to determine residual α_IIb_β_3_ activity and to diagnose aggregation-related defects. In the future, combining morphometric analysis with automated super-resolution microscopy could enable statistical analysis of platelet ultrastructure.

**Figure 6:**
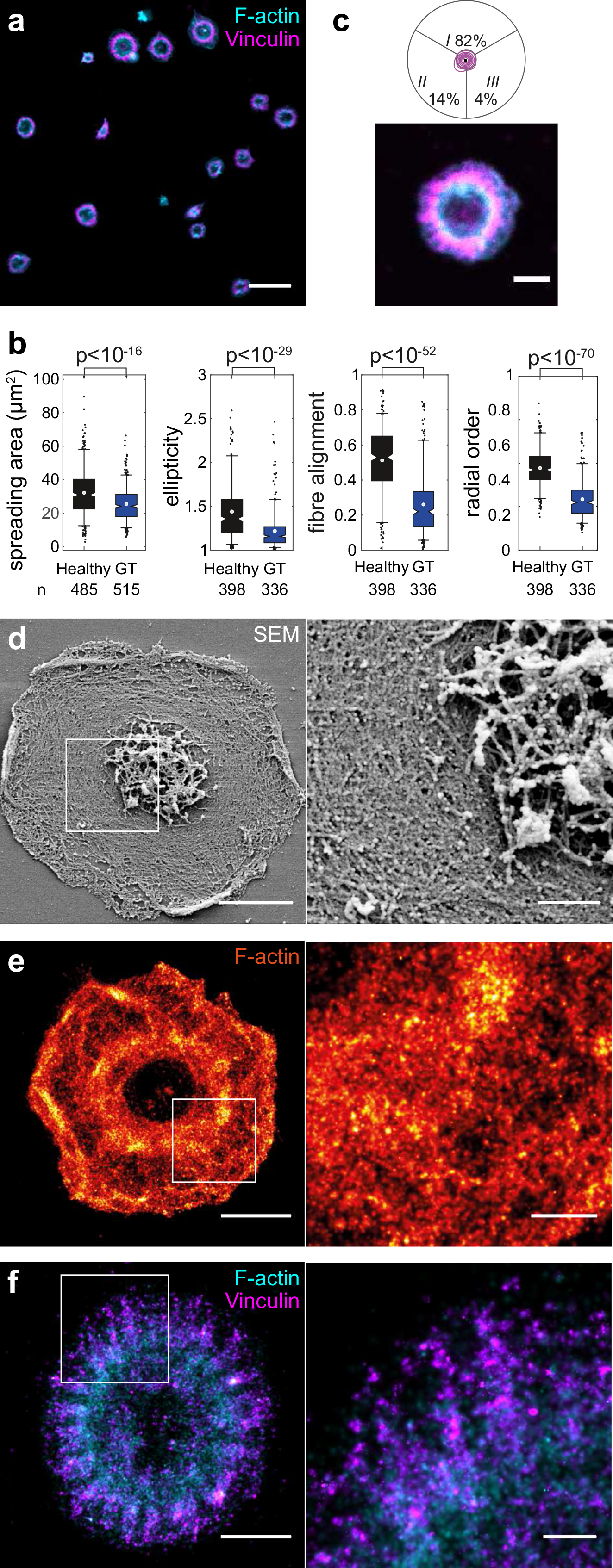
Cytoskeletal morphology on FG is altered in platelets from a patient with GT. a) Overview confocal micrograph with F-actin (cyan) and vinculin (magenta). b) Comparison of healthy and GT platelet morphology with respect to spreading area, ellipticity, fibre alignment, and radial order (cf. also Supplementary Figs. S19, S20, S24). c) Adhesion sites geometry and representative cell. d) Representative SEM image of detergent extracted GT platelet. e) dSTORM image of F-actin. f) Dual-colour dSTORM image of vinculin (magenta) and F-actin (cyan). Scale bars: 10 μm a), 2 μm (b and d-f, left column), 500 nm (d-f, insets). See also Supplementary Figs. S21 and S23 for overview dSTORM images and S16 for cryo-electron tomograms.

## METHODS

### Reagents

Reagents were purchased from Sigma Aldrich, if not mentioned otherwise. Acid citrate dextrose (ACD) tubes (Sol. B, Vacutainer®, BD, Switzerland); coverslips (18 mm diameter, thickness 1.5; Hecht-Assistent, Germany); human fibrinogen (FG;F3879); human fibronectin (FN; purified from plasma as described previously^60^); rat collagen type 1 (COL1; 354236, Corning, USA); murine laminin (LN; L2020); poly(acryl-amide)-g-(PMOXA, 1,6-hexanediamine, 3-aminopropyldimethylsilanol) (7000:4425:116.2:161.3 Mr; 0.2:0.4:0.4 d) (PAcrAm-g-PMOXA) and Poly(L-lysine)-graft-(poly(ethylene glycol)) (20'000:2000 Mr; 0.29 d) (PLL-g-PEG; gift from SuSoS AG, Switzerland); bovine serum albumin (BSA; 05470); Adenosine 5′-diphosphate sodium salt (ADP; A2754); eptifibatide (Integrillin; GlaxoSmithKline AG, U.K.); RUC-4 (gift from Prof. B.S. Coller, New York University); SiR-actin kit (CY-SC001, Spirochrome, Switzerland); monoclonal mouse anti-vinculin (V9131); monoclonal mouse anti-α-actinin (A5044); polyclonal rabbit anti-myosin IIa (3403, Cell Signalling Technology, USA); rabbit anti-phospho-myosin light chain 2 (3671S, Cell Signalling Technology, USA); goat anti-mouse Alexa Fluor 555 (A21424, ThermoFisher, USA); unconjugated donkey anti-mouse or anti-rabbit IgG (Jackson Immunoresearch, USA); Alexa Fluor 647 NHS ester (A37573, ThermoFisher, USA); CF680 NHS ester (92139, Biotium, USA); Alexa Fluor 488 Phalloidin (A12379, ThermoFisher, USA); Alexa Fluor 647 Phalloidin (A22287, ThermoFisher, USA); ProLong Gold Antifade Mountant (ThermoFisher, USA); 3-amino propyl tri-ethoxy silane (APTES;A3648); ammonium persulfate (APS;215589); tetramethylethylenediamine (TEMED;T7024); NHS-diazirine (26167, Thermo Fisher, USA).

### Blood collection and platelet isolation

This research was conducted according to the principles of the Declaration of Helsinki with approval by the Kantonale Ethikkommission Zürich (KEK-ZH-Nr. 2012-0111 and KEK-ZH-Nr. 2013-0027, respectively). All experiments were performed in accordance with these regulations, and informed written consent was obtained from all participants. Whole blood from healthy volunteers or from the GT patient was collected in ACD tubes at the University Hospital Zurich. Purification of platelets was performed not later than four hours after blood withdrawal, if not mentioned otherwise. 6 mL blood was gently transferred into 15 mL Falcon tubes and centrifuged at 180 g for 15 minutes at RT to obtain platelet-rich plasma (PRP). 1 mL of PRP was transferred into another Falcon tube without disturbing the buffy coat layer and spun at 900 g for 5 min. The platelet-poor plasma (PPP) was carefully pipetted away and the platelet pellet was gently resuspended in 400 μL Tyrode’s buffer (TB: 134 mM NaCl, 12 mM NaHCO_3_, 2.9 mM KCl, 0.34 mM Na_2_HPO_4_, 1 mM MgCl_2_, 10 mM Hepes, pH 7.4) at 37°C.

### Surface coatings

Coverslips were cleaned with piranha solution (1:1), washed with ddH_2_0, and blown dry under an airstream. Under standard conditions, surfaces were coated with human FG (50 μg ml^−1^ in PBS) for 1 h at room temperature (RT). Alternatively, FN, COL1 or LN were used accordingly. For surface ligand titration, the FG bulk concentration was varied while keeping the other parameters constant. Subsequent surface blocking was done by incubation with PAcrAm-g-PMOXA (100 μg ml^−1^ in PBS) for an additional hour at RT. Alternatively, PLL-g-PEG or BSA were tested as non-fouling agents, with inferior results.

### Preparation of poly-acrylamide (PAA) hydrogel substrates

Air-plasma treated glass coverslips (50mm diameter, #1.5) were incubated with APTES at RT for 3 min, washed thoroughly with water, incubated for 30 min with 0.5% (v/v) glutaraldehyde in PBS, washed again, and blown dry. 10 ml pre-gel solutions were prepared in different ratios of acrylamide and bis-acrylamide according to Tse and Engler^61^, degassed for 30 minutes, and mixed with 100 μl of 10% (w/v) APS, and 20 μl of TEMED, to initiate gelation. 3.9 μl of the activated gel solution were immediately pipetted onto an activated coverslip and a second glass coverslip (10mm diameter) was carefully placed on top. This sandwich was let to polymerize for 30 minutes and then the top coverslip was detached by using a scalpel and fine forceps. To covalently attach fibrinogen to the hydrogel surface, gels were covered with 20 μl of freshly dissolved NHS-diazirine (1 mg mL^−1^ in HEPES pH 8.2) and illuminated with UV (365nm, 0.125 W/cm^2^) for 2 minutes. After four quick repetitions of this step, samples were washed in HEPES, the buffer carefully removed and 20 μl FG solution (100 μg mL^−1^ in HEPES pH 8.2) were added onto the gels for 3 hours. Gels were then washed three times with HEPES and then equilibrated in Tyrode’s buffer for at least one hour. Seeding of platelets and fluorescence staining were performed as with the normal coverslips but without mounting.

### Sample preparation and fixation

Coated coverslips were placed in 12-well plates. Under standard conditions, 25 μl platelet solution was added to 500 μL pre-warmed TB containing 1 mM Ca^2+^ and 5 μM ADP and seeded on the coated coverslips at 37°C for 1 h. For titration experiments with Eptifibatide or RUC-4, the inhibitors were added to the buffer before the addition of platelets. After incubation, platelets were rinsed once with TB, detergent extracted with 0.25% (v/v) Triton X-100 and 3% (w/v) formaldehyde (FA) in cytoskeleton buffer (CB: 10 mM MES, 150 mM NaCl, 5 mM EGTA, 5 mM glucose, 5 mM MgCl_2_, pH 6.1) for 90 seconds, and subsequently fixed with 3% (w/v) FA in CB for 15 minutes at RT. For fixation of microtubules, 0.3% (w/v) and 2% (w/v) glutaraldehyde (GA) were used instead of FA during extraction and fixation, respectively, and samples were subsequently washed once and quenched with 0.1% (w/v) NaBH_4_ in PBS for 7 minutes. Samples for SEM were rinsed once with TB and detergent extracted with 0.75% (v/v) Triton X-100 in PHEM-buffer (60 mM Pipes, 25 mM Hepes, 10 mM EGTA, 2 mM MgCl_2_, pH 7.4) for two minutes, then fixed with 1% (w/v) GA in PHEM for 10 minutes at RT. After three rinses with PBS, all samples were stored at 4°C in PBS containing 0.02% (w/v) sodium azide.

### Fluorescence stainings

For (immuno)fluorescence, samples were permeabilized with 0.1% (v/v) Triton X-100 and 0.5% (w/v) BSA in PBS for 10 minutes and then blocked for additional 10 minutes with 3% (w/v) BSA in PBS at RT. For confocal microscopy, primary antibodies were diluted 1:80 in PBS containing 3% (w/v) BSA and incubated for one hour at RT, followed by three washes with PBS. Secondary antibodies (1:80) and phalloidin against F-actin (1:50) were incubated for another hour at RT, followed by three washes with PBS. Typically, phalloidin was in the 488 nm channel and the antibody in the 555 nm channel. Samples were mounted by ProLong Gold on 24 x 50 mm glass coverslips, let polymerize for 1 day at RT in the dark, and stored up to 4 weeks at 4°C.

For dSTORM, this procedure was modified as follows. Secondary antibodies were custom-labelled by NHS-reactive AF647 or CF680 yielding 1-2 dyes per antibody on average. Primary and secondary antibody concentrations were elevated (1:40), labelled samples were post-fixed with 3% (w/v) FA in PBS for 10 min, then washed and stored up to three days at 4°C in PBS. For d-STORM imaging of F-actin, phalloidin-AF647 (1:20) was incubated for 30 min, followed by three quick washes in PBS and imaged immediately afterwards. Alternatively, phalloidin-AF488 was used as a co-stain for epifluorescence. For dual-colour dSTORM, AF647 and CF680 labels were used.

### Optical microscopy

Mounted samples were imaged on a Leica SP5 laser scanning confocal microscope (Leica Microsystems, Germany) using a 63x oil immersion objective and 2x zoom (resulting in a field of view of 123 x 123 μm at 60.1 nm pixel size) and excitation at 488 nm and 546 nm. For statistical analysis of platelet morphology, between 200-250 cells were recorded per condition at 6-8 different field of views on the sample. Samples for dSTORM were transferred to a custom holder with imaging buffer (200 mM Tris, pH 8.2, 4% (w/v) glucose, 1 mg ml^−1^ glucose oxidase, 0.2 mg ml^−1^ catalase, 20 mM TCEP, 2.5% glycerol and 35 mM cysteamine). dSTORM imaging was carried out on a home-built setup as described previously^60^. Fitting and analysis of dSTORM movies was performed using the software package SMAP (courtesy of Dr. Jonas Ries, EMBL Heidelberg).

### Electron microscopy

For SEM, further fixation with osmium tetroxide (OsO_4_, 0.5% (w/v) in ddH2O) for one hour was followed by dehydration in a graded series of ethanol (from 50% to 100% in ddH2O). Subsequently, the samples were dried over the critical point of CO_2_ (T_c_: 31°C, P_c_: 73.8 bar) using a critical-point dryer (Tousimis CDP 931, USA). After sputter coating with 5 nm platinum (MED010, Balzers, Liechtenstein) images were recorded with a Zeiss Leo-1530 scanning electron microscope at 5 kV acceleration voltage detecting secondary electron signals. For cryo-EM, please refer to the Supplementary Information.

### Image analysis

A detailed description of the image analysis is given in the Supplementary information. In short, outlines and morphological features of single platelets were automatically extracted from fluorescence images by different morphological and filtering operations. The outline yielded the single cell spreading area and ellipticity, i.e. the ratio of the long to the short axis of an equivalent ellipse. The F-actin alignment per cell was quantified by the average of the cosine of the local orientation of actin fibres relative to their mean orientation^22^. The radial order per cell was quantified in an analogous fashion but relative to the local position of the fibres from the centroid. The radial distribution profile of the vinculin staining was subjected to a Fourier decomposition and the resultant amplitudes were used to assign a characteristic morphology (isotropic, bipolar, triangular) to the cell. Then, the distribution of many platelet morphologies was depicted as a contour plot.

### Statistics

A comprehensive description of the statistical analysis is found in the Supplementary information. Briefly, boxes in boxplots represent upper and lower quartiles, notches depict the median and comparison intervals, the small circle marks the mean. Whiskers represent the 5^th^ and 95^th^ percentiles, and outside data are depicted as dots. A non-parametric Kruskal-Wallis rank test with Scheffe post-hoc testing was applied to make (multiple) comparisons as some parameters did not follow a normal distribution. As a general remark, Kruskal-Wallis was always less discriminative than ANOVA with Tukey-Kramer. The p-values for comparisons are reported and assumed to be significant for p < 10^−4^ (n.s.: not significant).

### Data availability

The datasets generated during and/or analysed during the current study are available from the corresponding author on reasonable request.

## Acknowledgements

We thank the GT patient M.T. for her consent and her time to participate in this study. We thank Prof. Barry S. Coller for insightful discussions and for generously providing the RUC compounds. We gratefully acknowledge Dr. Stefan Zürcher (SuSoS AG) for the generous gift of the non-fouling agents PAcrAm-g-PMOXA and PLL-g-PEG. We thank Prof. J. Oldenburg and Dr. A Pavlova (Institute for Experimental Hematology, University Hospital Bonn) for performing the analysis of ITGA2B and ITGB3 genes. We would like to acknowledge Tanuj Sapra for helpful discussions and his initiative during the initial stage of the project, Dr. Jens Moeller and Dr. Lina Aires for assistance with hydrogels, Lukas Braun for preparing Supplementary Fig. S17, Dr. Isabel Gerber for assistance with EM preparation, and Dr. Tobias Schwarz and Joachim Hehl (ScopeM, ETH Zurich) for assistance with STED and SIM microscopes. We thank Dr. Jonas Ries (EMBL Heidelberg) for kindly providing software for the visualization of dSTORM data. This work was supported by a grant from the Swiss Science Foundation (CR32I3_156931; V.V.), by the Wyss Zurich Translational Center (2-72187-16; V.V.), by ETH Zurich, and by RCSI (I.S.)

## Author contributions

I.S., V.V. and O.M. laid out the concept of the study; S.L. and I.S. planned experiments; S.L. performed all platelet spreading experiments and confocal microscopy; I.S. performed livecell imaging; S.L. and I.S. performed super-resolution imaging; S.S. performed cryo-ET; S.L. performed SEM; I.S. developed the image analysis scripts; S.L. prepared hydrogels; S.L. and I.S. analysed the data; J.D.S. recruited and characterized the GT patient; S.L. and I.S. drafted the manuscript; and all authors contributed to the interpretation of data and critically revised the report.

## Additional information

ORCID profiles: I.S., 0000-0002-5699-1160; V.V., 0000-0003-2898-7671.

